# Allometric equations for estimating tree volume and aboveground biomass of *Acacia mangium* Willd. in the Batéké Plateau, Republic of Congo

**DOI:** 10.1101/2025.10.15.682680

**Authors:** Chelton Desarmes, Grace Jopaul Loubota Panzou, Flore Hirsch, Paul Bertaux, Nicolas Bayol

## Abstract

*Acacia mangium* Willd. is one of the most widely planted fast-growing tropical species, yet it remains poorly documented in Central Africa, particularly in terms of biomass accumulation. This knowledge gap, combined with the lack of robust allometric equations, limits the development of reliable Measurement, Reporting, and Verification (MRV) protocols for plantations. Since accurate estimates of tree volume and aboveground biomass (AGB) are essential for carbon stock assessment and for implementing climate change mitigation strategies such as REDD+, this study developed allometric equations for tree volume and AGB in *A. mangium* plantations on the Batéké Plateau, Republic of Congo, using destructive sampling of 54 trees (4–24 cm DBH; 25, 42, and 66 months old). Trees were measured, felled, and partitioned into trunks and large branches (>4 cm), small branches (1–4 cm), foliage, and stumps. Samples of each compartment were oven-dried and weighed. Results showed that AGB allocation to trunks and large branches increased with age, while allocation to small branches declined. A power-law model (Y = a(D^2^H)^b^) provided the best fits for both volume (Y= 0.401(D^2^H)^0.984^) and AGB (Y = 0.259(D^2^H)^0.900^). When applied to inventory data from permanent sample plots in plantations in the Batéké Plateau, estimates rose from 38.3 m^3^ ha^−1^ and 29.4 Mg ha^−1^ at 28 months to 157.6 m^3^ ha^−1^ and 105.5 Mg ha^−1^ at 64 months. These equations provide a robust reference for *A. mangium* in the Batéké Plateau and will strengthen the accuracy of biomass and carbon monitoring for effective forest-based climate mitigation initiatives.

## Introduction

Estimating aboveground tree biomass to assess carbon stocks is essential for quantifying the carbon balance in tropical forests. Such estimation has become a global priority, particularly for the implementation of initiatives such as Reducing Emissions from Deforestation and Forest Degradation (REDD+) and Afforestation, Reforestation and Revegetation (ARR) [1,2]. The successful implementation of such initiatives in tropical countries plays a key role in global climate change mitigation, and participating countries must provide detailed strategies and carbon reference levels [3]. However, establishing these reference levels requires accurate estimates of biomass and carbon stocks.

Traditionally, allometric relationships have been used to estimate stem volume or tree biomass [4]. Allometry describes the relationship between a tree’s measurable characteristics, such as diameter and/or height, and another variable that is more challenging to measure directly, such as biomass or volume [5,6]. A major limitation of allometric models is that their initial development requires the destructive sampling of trees [7]. Furthermore, selecting the most appropriate model is essential to improve accuracy [8], yet identifying the best-performing equation is not always straightforward [9].

While volume estimation is critical for forest management and timber commercialization, the demand for biomass estimation has significantly increased over the past decade due to its relevance in climate change mitigation [10]. In the Congo Basin countries, the establishment of robust Measurement, Reporting, and Verification (MRV) protocols is hindered by the lack of data necessary for developing allometric equations. This data gap is particularly critical for plantations, which remain among the most effective strategies for mitigating climate change [11]. Developing region-specific equations can significantly improve biomass estimation and MRV protocols [7].

In the Republic of Congo, agroforestry and forestry plantations play a key role in reforestation programs, where they are at the heart of ecological restoration and carbon sequestration strategies [12,13]. They also contribute substantially to wood and energy production, meeting the rapidly growing urban demand for firewood and charcoal [14,15]. In 2013, the government of the Republic of Congo took a proactive approach to afforestation and reforestation, notably through the National Afforestation and Reforestation Program (PRONAR), which aimed to establish 1 million hectares of forest and agroforestry plantations [15–17]. Among the species used, *Acacia mangium* is particularly prominent due to its rapid growth, tolerance of nutrient-poor and acidic soils, and ability to fix atmospheric nitrogen through symbiotic associations with *Rhizobium* bacteria, making it particularly suitable for the long-term success of afforestation initiatives [13,18–22]. In addition, it stands out for its substantial potential in terms of timber and energy production, making it even more attractive for planting programs in tropical areas [13,23,24].

In this context, this study aims to develop and test allometric equations adapted to *Acacia mangium* plantations for tree volume and aboveground biomass on the Batéké Plateau, Republic of Congo. This study has two specific objectives to: i) develop allometric equations for estimating wood volume and aboveground biomass of trees; and ii) Assess the impact of applying allometric equations for volume and AGB to these plantations. This study will enable significant advances in several fields: forest inventory, carbon stock assessment and sustainable management of *Acacia mangium* plantations on the Batéké Plateau and overall the Central Africa.

## Materials and methods

### Study sites

This study was conducted in the plantation of *Acacia mangium* Willd. (Fabaceae) located on the Batéké Plateau in the Republic of Congo. The Batéké Plateau of the Republic of Congo consists of four distinct savannah sub-plateaus : Koukouya, Djambala, Nsa-Ngo and Mbé/Batéké [25,26]. Fieldwork for this study was carried out in three sites representing three main topography types in the Batéké Plateau landscape. The first site (Site 1: 3º52’2’’S, 15º31’5’’E) was located on the Mbé plateau with a stable topography. The second site (Site 2: 3º37’44’’S, 15º20’9’’E) was assigned to a transition area, with an undulating topography, between the Mbé plateau and the hills. The third site (Site 3: 2º39’27’’S, 15º51’6’’E) has been described as an area with a succession of hills (Figure 1). All three sites are characterized by tropical transitional climate with an average annual rainfall of 1500–1800 mm. There is a main dry season from June to September and a shorter one in January and February. The average annual temperature is 25°C. The soils are predominantly ferrallitic, sandy with low clay content, leached with low organic matter and mineral content, classifying them as low fertility soils [25,26].

**Figure 1.**
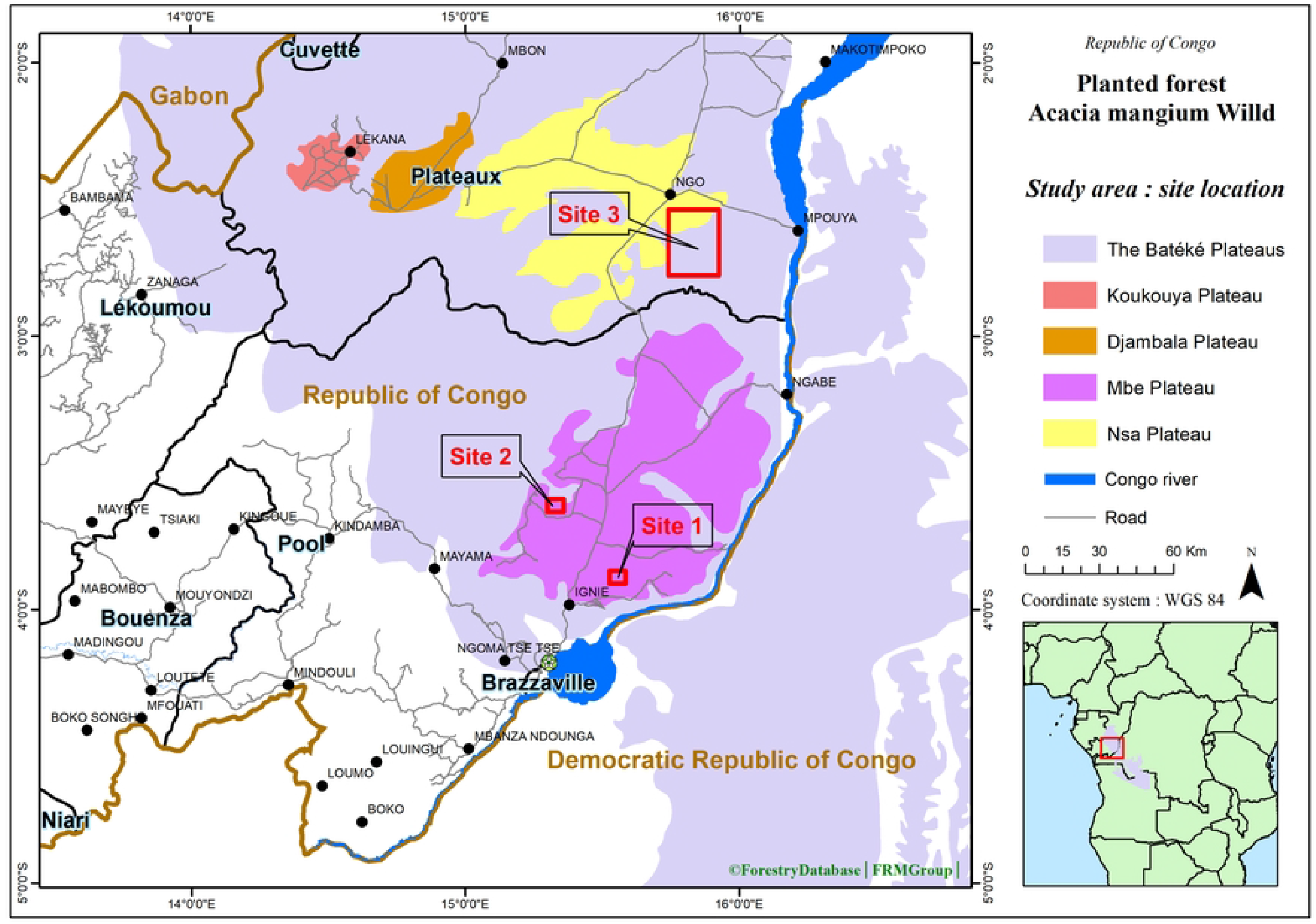
Location of study sites on the Batéké Plateau in Republic of Congo.

### Experiment design and tree sampling

The experimental design was based on pre-existing forest inventory data, comprising 2,722 measurements of diameter at breast height (DBH) collected across the three sites with trees aged 25, 42, and 66 months old. Specifically, Site 1 (66 months old) contributed 85 DBH measurements, Site 2 (25 and 42 months old) contributed 640 measurements, and Site 3 (25 months old) contributed 1,997 measurements. DBH values ranged from 2 cm to 23 cm. Site 3 was subdivided into two sub-units due to pronounced differences in growth observed between two plantation zones, despite their identical age.

The pre-existing tree DBH measurements data were divided into four classes corresponding to the quartiles of the stand’s DBH distribution (Supporting Information Figure S1). DBH values below 4 cm were excluded from the sampling frame, as they are younger trees planted to replace dead ones and should have specific allometric equation [27]. The sampling intensity in term of number of tree for each class was proportionally adjusted on the standard deviation of basal area [6]. A total of 54 trees from all diameter classes and plantation ages were selected in all three sites (Table 1). This number of sampled trees exceeds the recommended minimum (30) for homogeneous plantation stands [6,28,29] and should ensure sufficient precision for the resulting models.

**Table 1.**
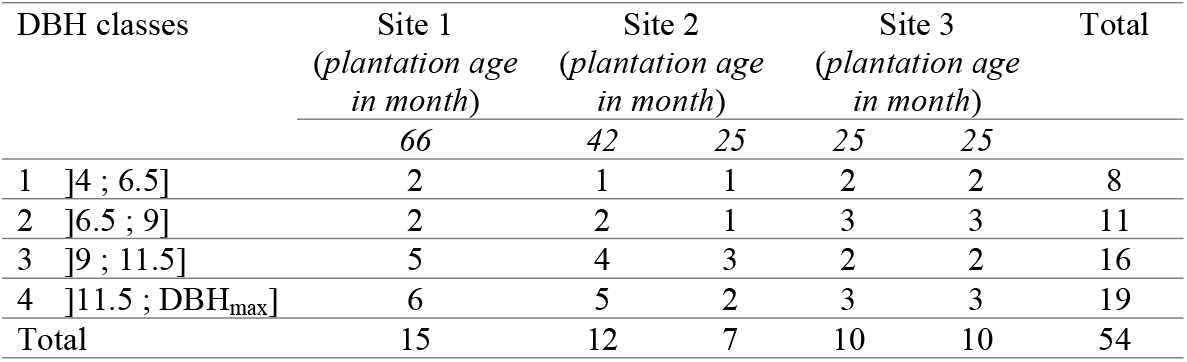
Number of trees according to DBH class and plantation age in each site.

### Tree measurements

Prior to felling, non-destructive tree diameter and height measurements were performed in 2024. The diameter at breast height (DBH) was measured at 1.30 m above the ground using a diameter tape (Richter model: 283D/5) for single-stem trees and multi-stemmed trees (number of stems ranged from 1 to 6). For multi-stemmed tree, the diameter was calculated as the root mean square of stems diameter [30]. Each stem with a DBH of at least 1 cm at breast height was measured. For each standing tree, total height was measured three times from three different positions using a SUUNTO clinometer (SUUNTO model: PM-5/1520 PC), ensuring that the operator stands at a minimum distance equivalent to the height of the tree to be measured.

Destructive sampling was carried out using a chainsaw. Each tree was cut down at 10 cm from the base. The length of felled tree was measured before being cut into four compartments: trunks and large branches (> 4 cm diameter), small branches (1-4 cm diameter), foliage and stump. The trunks and large branches were cut into chunks less than 2 m long [6]. The length and diameter (butt and butt end) of each chunk were measured for each felled tree.

For each compartment, the fresh mass was measured in the field using a 500g precision mechanical balance. Representative samples (trunk: discs of 5 cm at both ends and every 2 m; branches divided into 4 diameter classes: on average, 5 fixed-length samples per class; phyllodes: 3 bags of 10 liters each) were collected and weighed on a 1 g precision balance. These samples were then dried in a circulating air oven at 105°C (for stems, large branches and small branches) or 70°C (for foliage) until their mass stabilized [6,31]. The dry mass of each representative sample was measured using the 1 g precision balance.

For each tree, the observed volume (*V*_*obs*_) was calculated as the sum of the chunks and stump volumes. The volume of each chunk was calculated using Smalian’s formula [32] and the volume of the stump using the truncated cone formula [33]. The observed dry aboveground biomass (*AGB*_*obs*_) was calculated as the sum of the dry aboveground biomass of all the compartments. The dry biomass of each compartment, except the stump, was calculated by multiplying the fresh mass of the compartment by the dry mass/fresh mass ratio determined from the dried samples. The stump biomass was estimated from its volume using the relationship: *AGB*_*stump*_ *= V*_*stump*_ *\* d*, where d is the anhydrous density (0.507 g/cm^3^) of *Acacia mangium* [34].

### Data analysis

To develop the allometric equations, both non-destructive measurements (diameter at breast height, *D*, and total height, *H*) and the destructive measurements (observed volume, *V*_*obs*_, and aboveground biomass, *AGB*_*obs*_) were used. Allometric models using D as a predictor have been widely used in allometric studies due to their simplicity and operational applicability [31,13,7,35]. However, integrating total height with diameter (*D*^*2*^*H*) in the allometric models can significantly improve the accuracy of AGB estimates [8,36–38]. Since tree height measurements are difficult in tropical forest [39,40], we first evaluated how accurate are non-destructive tree height measurements in comparison with destructive measurements on felled trees. Given the strong correlation (Pearson’s r = 0.98, p-value < 0.001) and no significant difference between the non-destructive and destructive height measurements (p-value > 0.05; repeated measures analysis of variance; see supporting information Table S1, and Figure S2), the non-destructive height measurement is confidently used to establish volume and AGB allometric equations.

For each predictor (D in cm or D^2^H in m^3^), four allometric models (linear, power, exponential and logarithmic) commonly used worldwide to estimate volume or aboveground biomass were tested [35]. The modelling approach used the following equation based on the diameter (D) and total height (H) of a tree *i* belonging to all three sites:

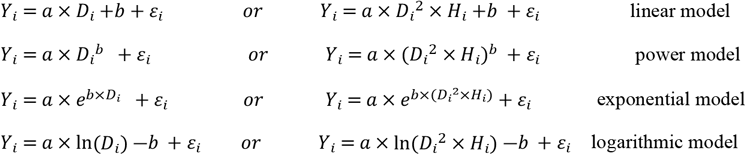

where a and b are the fitted coefficient or exponent, *Y*_*i*_ is alternatively the observed tree volume (V_obs_ in m3) or the aboveground biomass (AGB_obs_ in Mg) and *ε*_*i*_ is the error.

For the power and exponential models, initial parameter values were derived from linear regressions on log-transformed data. These starting values were then used to fit the final models directly to the untransformed data using the “nls()” function, thereby avoiding correction of retransformation bias [41,42].

The biomass expansion factor (BEF) which represents the part of biomass contained in the small branches (D < 4 cm) and phyllodes is useful to convert tree volume to aboveground biomass [43]. It was obtained from the linear regression: *AGB*_*obs*_ *= BEF* (*V_obs *_ *d*), where V_obs_ is of the tree volume, and *d* the dry density (0.507) of *A. mangium* [34].

Model selection was based on the following criteria: (a) the coefficient of determination (R^2^), representing the proportion of explained variance; (b) model performance indicators, including root mean square error (RMSE) and prediction bias; (c) Akaike’s Information Criterion (AIC), and Bayesian Information Criterion (BIC) which balances model accuracy and complexity. A lower AIC or BIC value indicates a better model fit [44]. To estimate the predictive performance of the selected models, a leave-one-out cross validation (LOO-CV) was computed and evaluate the mean absolute error (MAE), the mean relative error (MRE), and the mean bias of the models. The LOO-CV compares the model’s predictions with independent variables that were not used during model fitting [45–47].

To assess the impact of volume and AGB allometric equations developed in this study, forest inventory data were converted into volume and AGB using the best fitted models. The forest inventory data consisted of eight permanent sample plots (PSP), with two PSP established in Site 1 (64 months old) and six in Site 2 (40 months old for 4 PSP and 28 months old for 2 PSP). Each PSP corresponds to three lines of thirty spots, i.e. approximately 0.0907 ha. The tree density was estimated at approximately 715 trees/ha in the Site 1 and 880 trees/ha in the Site 2. The forest inventory data (DBH and H) for all living trees (a total of 634 trees) in the PSP was collected in March 2024. These forest inventory data were used to assess the impact of these equations for plantation ages ranging from 25 to 66 months. A linear regression was fitted between the estimated volume or AGB, and plantation age to assess the validity of allometric equations across the plantation age range (25-66 months).

All statistical analyses were computed using the open-source R environment (R version 4.4.2, R Core Team, 2024).

## Results

### Overview of sampled trees

The 54 sampled trees have a mean diameter at breast height (DBH) of 11.0 ± 4.5 cm and a mean height (H) of 10.3 ± 3.8 m (Table 2). DBH and H ranged from 4.5 to 23.8 cm and 5.3 to 20.2 m respectively, indicating the range of validity of the allometric models. The observed stem volume and aboveground biomass (AGB) of felled trees ranged from 0.007 to 0.477 m^3^ (mean = 0.073 ± 0.091 m^3^) and from 0.004 to 0.284 Mg (mean = 0.052 ± 0.059 Mg), respectively (Table 2).

**Table 2.** Mean ± standard deviation of the tree measurements (DBH, total height, volume, and AGB) in each study site.

| Site | Plantation age (month) | Mean DBH (cm) | Mean height (m) | Mean volume of felled tree (m<sup>3</sup>) | Mean AGB of felled tree (Mg) |
| --- | --- | --- | --- | --- | --- |
| Site 1 | 66 | $12.7 \pm 5.7$ | $14.6 \pm 3.8$ | $0.135 \pm 0.142$ | $0.089 \pm 0.093$ |
| Site 2 | 42 | $12.0 \pm 4.4$ | $10.3 \pm 1.6$ | $0.041 \pm 0.027$ | $0.032 \pm 0.021$ |
| | 25 | $9.9 \pm 2.8$ | $8.4 \pm 1.4$ | $0.070 \pm 0.055$ | $0.050 \pm 0.036$ |
| Site 3 | 25 | $9.7 \pm 4.4$ | $9.2 \pm 1.8$ | $0.048 \pm 0.051$ | $0.038 \pm 0.039$ |
| | 25 | $9.5 \pm 3.3$ | $6.3 \pm 1.1$ | $0.030 \pm 0.027$ | $0.028 \pm 0.023$ |
| mean | 44 | $11.0 \pm 4.5$ | $10.3 \pm 3.8$ | $0.073 \pm 0.091$ | $0.052 \pm 0.059$ |

The allocation of tree AGB varied with plantation age (Figure 2). The proportion of AGB allocation increased for the trunk and large branches and decreased for small branches and phyllodes according to plantation age from 25 months to 66 months.

**Figure 2.**
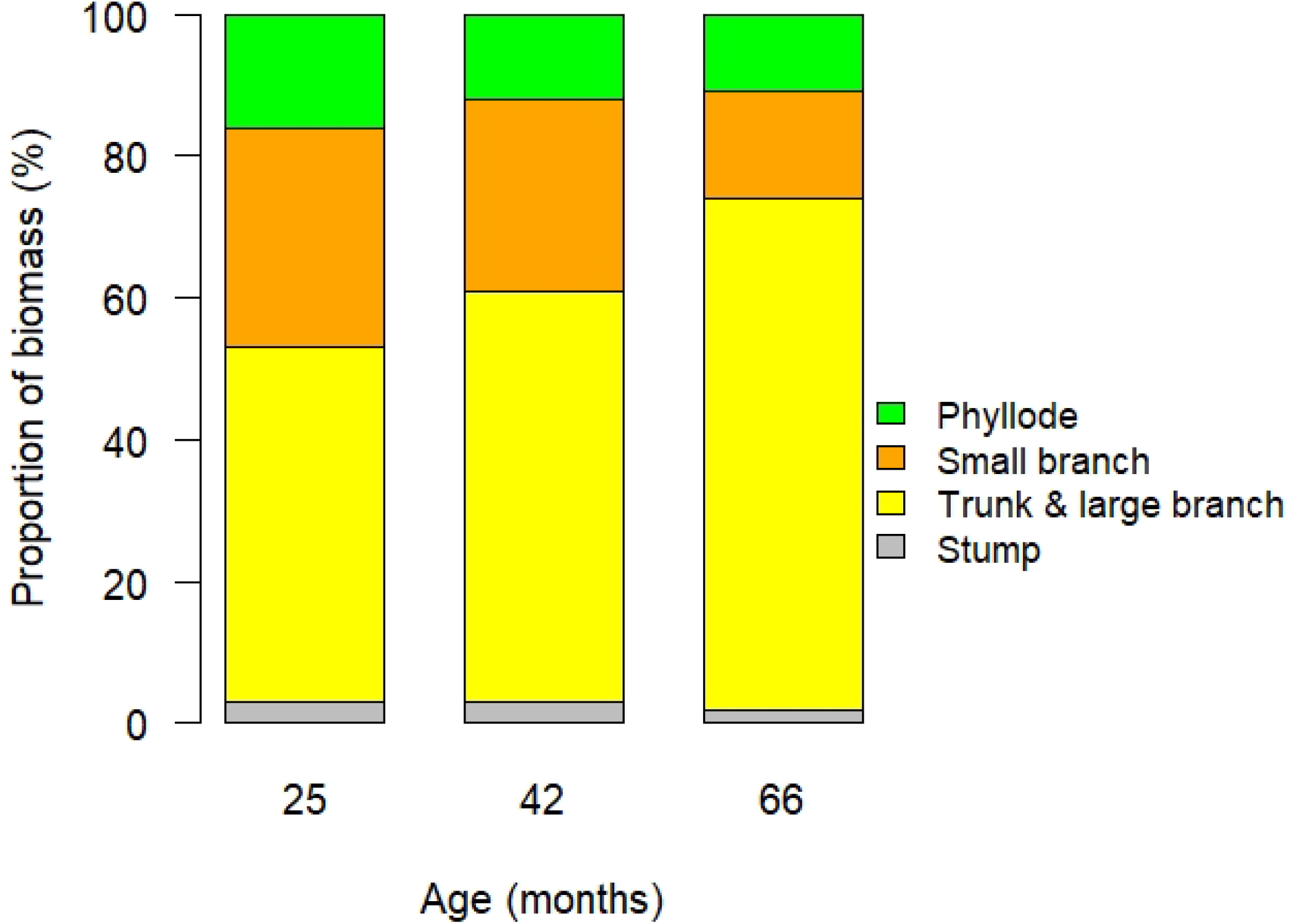
Distribution of the AGB allocation according to the compartments (Trunk and large branch, small branch, phyllode and Stump) and plantation ages.

The proportion of biomass contained in the small branches (D < 4 cm) and phyllodes, which represents the biomass expansion factor (AGB = BEF*(V_obs_ \**d*)), varied significantly with stand age (Table 3). At 25 months, the BEF reached 1.57 ± 0.04 and progressively decreased with age, being 1.36 ± 0.06 at 42 months and 1.28 ± 0.05 at 66 months. Model fits were very strong, with coefficients of determination ranging from 0.977 to 0.983.

**Table 3.**
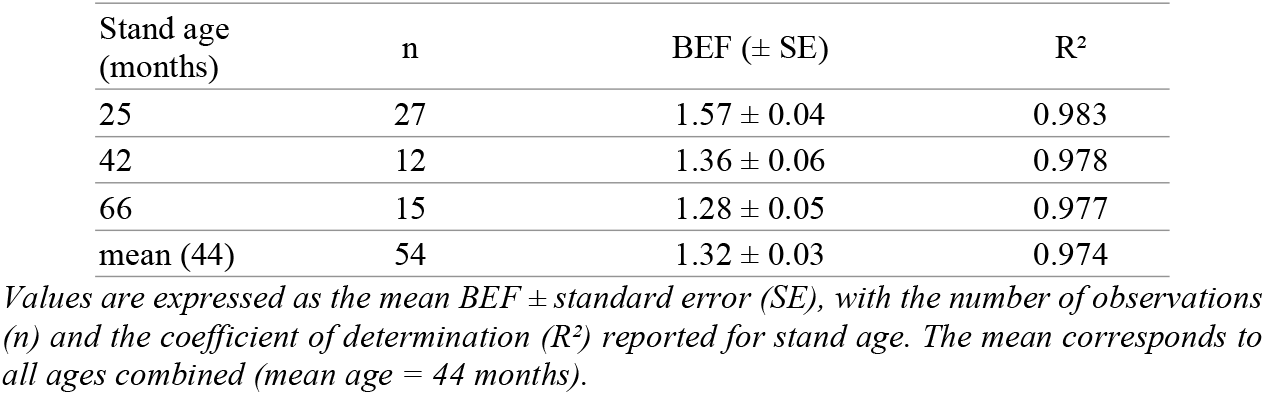
Biomass Expansion Factor (BEF) by stand age and for the global model.

When all ages were considered together, the global model (mean age of 44 months, n = 54) yielded a BEF of 1.32 ± 0.03 (R^2^ = 0.974). This overall value is close to that obtained at 42 months and well reflects the general trend, namely a progressive decline in BEF with increasing stand age.

### Allometric models for volume and aboveground biomass

Models using D^2^H as a predictor exhibited better performance than those using only D^2^, except for the exponential model (Table 4). These findings indicate that D^2^H provides a more accurate representation of the variability in observed tree volume and aboveground biomass. The power model with D^2^H as predictor (Model 4: *Y = a(D*^*2*^*H)*^*b*^) demonstrated the best overall performance for both volume and AGB allometric equations (Table 4).

**Table 4.** Allometric equations for tree volume (V) and aboveground biomass (AGB).

| Models | N° Equation | Coefficient and exponent |  | Performance criteria |  |  |  |  |
| --- | --- | --- | --- | --- | --- | --- | --- | --- |
|  |  | a | b | R<sup>2</sup> | AIC | BIC | RMSE | Bias |
| 1 | $V = aD + b$ | 0.01785 | -0.12377 | 0.79 | -184.40 | -178.43 | 0.041507 | 40.83 |
| | $AGB = aD + b$ | 0.01191 | -0.07889 | 0.83 | -243.14 | -237.17 | 0.024094 | 33.23 |
| 2 | $V = a(D^2H) + b$ | 0.40053 | 0.00167 | 0.98 | -310.89 | -304.92 | 0.012867 | -3.98 |
| | $AGB = a(D^2H) + b$ | 0.25869 | 0.00630 | 0.97 | -333.17 | -327.20 | 0.010469 | -12.48 |
| 3 | $V = aD^b$ | 0.00005 | 2.89288 | 0.92 | -234.45 | -228.49 | 0.026112 | 5.28 |
| | $AGB = aD^b$ | 0.00007 | 2.60588 | 0.93 | -291.51 | -285.54 | 0.015396 | 3.09 |
| 4 | $V = a(D^2H)^b$ | 0.40115 | 0.98435 | 0.98 | -310.80 | -304.83 | 0.012878 | -1.29 |
| | $AGB = a(D^2H)^b$ | 0.25917 | 0.90017 | 0.97 | -335.49 | -329.53 | 0.010245 | -3.66 |
| 5 | $V = ae^{bD}$ | 0.00753 | 0.17491 | 0.92 | -234.26 | -228.30 | 0.026158 | -33.43 |
| | $AGB = ae^{bD}$ | 0.00676 | 0.16063 | 0.92 | -283.48 | -277.51 | 0.016584 | -30.54 |
| 6 | $V = ae^{b(D^2H)}$ | 0.04470 | 2.18624 | 0.85 | -202.16 | -196.19 | 0.035214 | -102.42 |
| | $AGB = ae^{b(D^2H)}$ | 0.03350 | 2.04148 | 0.82 | -238.80 | -232.84 | 0.025081 | -87.28 |
| 7 | $V = a\ln(D) - b$ | 0.18079 | -0.34663 | 0.63 | -154.84 | -148.87 | 0.054576 | 42.87 |
| | $AGB = a\ln(D) - b$ | 0.12180 | -0.23032 | 0.68 | -208.92 | -202.95 | 0.033076 | 39.46 |
| 8 | $V = a\ln(D^2H) - b$ | 0.07286 | 0.24022 | 0.72 | -168.65 | -162.69 | 0.048022 | 56.12 |
| | $AGB = a\ln(D^2H) - b$ | 0.04797 | 0.16247 | 0.74 | -219.03 | -213.06 | 0.030120 | 37.69 |

For each model, the estimated coefficients (*a*, *b*) and performance criteria are reported, including coefficient of determination (*R*<sup>2</sup>), Akaike Information Criterion (*AIC*), Bayesian Information Criterion (*BIC*), root mean square error (*RMSE*), and bias. Volume is expressed in cubic meters (m<sup>3</sup>), AGB in megagrams (Mg), *D* in centimeters (cm), and *D*<sup>2</sup>*H* in cubic meters (m<sup>3</sup>).

For tree volume, Model 2 (linear) and Model 4 (power) showed comparable performance. Their respective goodness-of-fit statistics were: R^2^ = 0.98 for both models; AIC = −310.89 and −310.80; BIC = −304.92 and −304.83; RMSE = 0.012867 m^3^ and 0.012878 m^3^; and bias = −3.98 % and −1.29 %, respectively (Table 4).

Results from the leave-one-out cross-validation (LOOCV) indicated a mean relative error (MRE) of 13.45 % for the linear model (Model 2) and 12.28 % for the power model (Model 4). The mean absolute error (MAE) was nearly identical for both models (0.0080 m^3^ and 0.0081 m^3^, respectively). Prediction bias remained negligible, with values of −0.0001 m^3^ for Model 2 and −0.0005 m^3^ for Model 4.

For the aboveground biomass (AGB), the model 4 “*AGB = a(D*^*2*^*H)*^*b*^*”* exhibited the best performance with the highest R^2^ (0.97), the lowest AIC (−335.49) and BIC (−229.53), and the lowest RMSE (0.010245) and bias (−3.66). The LOOCV results further confirmed the robustness of model 4, with a MAE of 0.0076 Mg, an MRE of 19.08%, and a negligible mean bias of −0.0001 Mg, indicating good predictive accuracy at the individual tree level (Table 4).

### Application of volume and AGB allometric models

The power model (model 4) was chosen as the best model to estimate volume and AGB. Using data from the eight permanent sample plots from 28 to 64 months, our results showed a progressive increase in volume and AGB with plantation age (Figure 3). The volume increased from 38.29 m^3^ ha^−1^ at 28 months to 157.62 m^3^ ha^−1^ at 64 months, with a mean annual increment from 16.41 m^3^ ha^−1^ year^−1^ to 29.78 m^3^ ha^−1^ year^−1^. Similarly, AGB varied from 29.41 Mg ha^−1^ at 28 months to 105.51 Mg ha^−1^ at 64 months, with a mean annual increment from 12.60 Mg ha^−1^ year^−1^ to 19.94 Mg ha^−1^ year^−1^. This indicates that the plantation is in an ascending growth phase.

**Figure 3.**
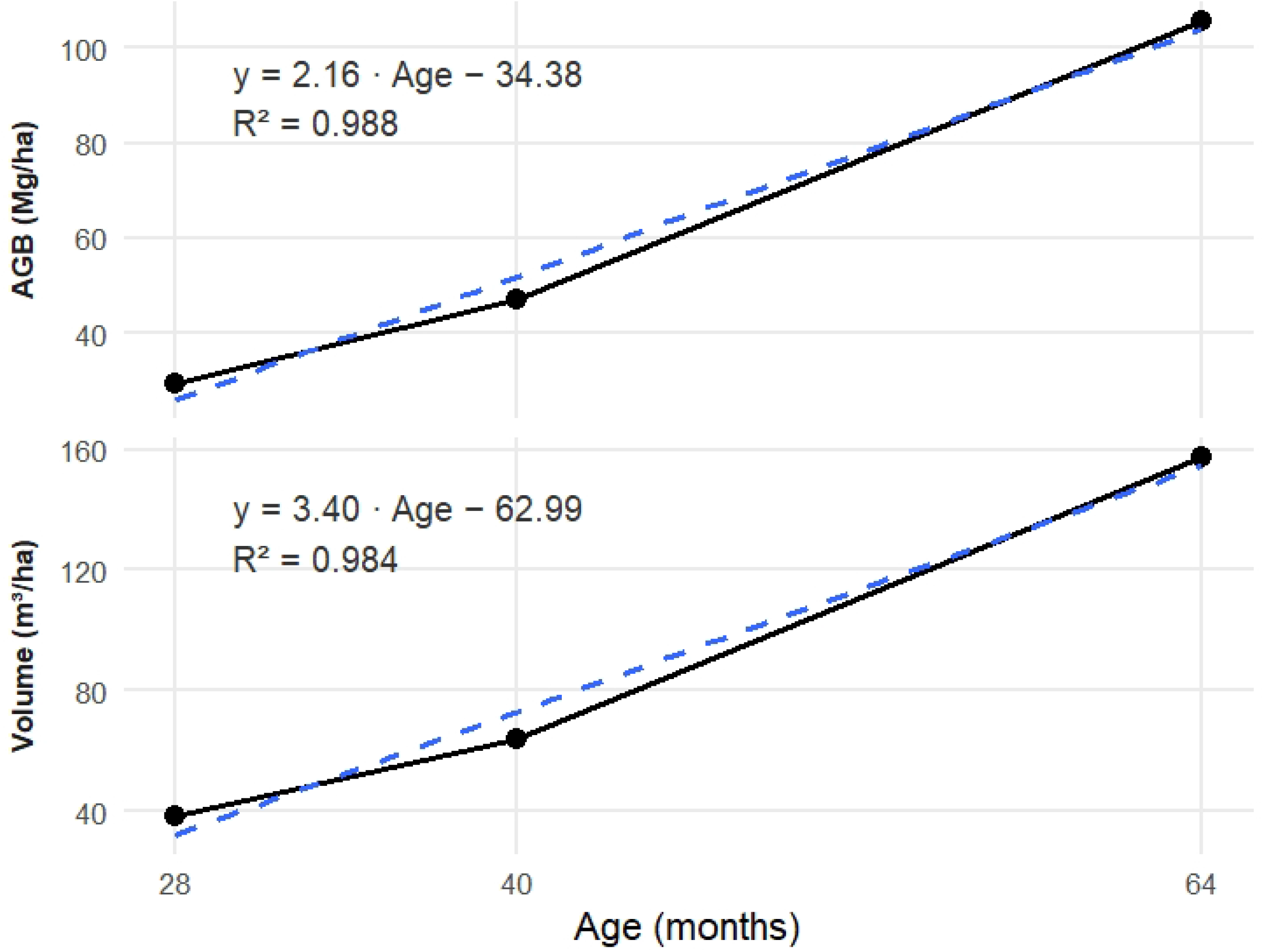
Volume and aboveground of 8 PSP located on Site 1 and 2

## Discussion

### Aboveground biomass partitioning

Destructive sampling revealed that the trunk and large branches accounted for the majority of aboveground biomass (> 50%) across all plantation ages, consistent with findings for *A. mangium* in Indonesia [43] and *A. mearnsii* in Ethiopia [48]. The increasing allocation to these compartments with age supports the relevance of the selected diameter range for validating allometric equations, although the ontogenetic scope remains limited (4–24 cm DBH) compared to natural forests (> 200 cm). Consequently, this study will need to be updated to account for larger diameter classes.

The Biomass Expansion Factor (BEF) estimated in this study for the 54 sampled trees aged 25 to 66 months was 1.32. This value is very close to the estimate of 1.332 reported by Miyakuni [43] for *Acacia mangium* trees aged 36 to 96 months (3–8 years), suggesting a relative stability of BEF within this age range. However, both values are significantly lower than the default BEF value of 3.4 (range: 2.0–9.0) recommended by the Intergovernmental Panel on Climate Change (IPCC) guidelines for tropical forests, which includes *Acacia mangium* [49].

BEF is known to vary with multiple factors, including forest type, tree age, stand density, growth conditions, and climate [36]. It has been shown in this study, in agreement with the findings of Miyakuni [43], that BEF decreases with stand age. The use of age-class-specific BEF values is therefore regarded as a relevant approach to improve the accuracy of biomass estimates and to reduce uncertainties in carbon stock assessments. Nevertheless, BEF does not conform to any classical allometric model. As highlighted by Miyakuni [43] and further supported by the present study, no statistically robust relationships could be established between BEF and commonly used predictors such as D, H, or D^2^H (see Figure S3 in Supporting Information). This lack of predictive consistency underscores the empirical nature of BEF and suggests that its estimation should rely on direct measurements rather than on allometric surrogates.

### Allometric equations for volume and aboveground biomass estimation

The choice of the allometric model is a critical step in the estimation of forest aboveground biomass [50,51]. As demonstrated in this study, several studies reported that the inclusion of multiple predictors in combination with tree diameter, provides a better estimate of tree volume and aboveground biomass [6,52]. The power model with D^2^H as predictor showed the best overall performance for the volume and AGB allometric equations. The leave-one-out cross-validation (LOOCV) results also showed a better predictive performance for the power model in terms of relative accuracy. Previous studies have also shown that power functions were able to estimate AGB with a high goodness of fit [53]. It is the most commonly used functional form in allometric equations for estimating the tree volume and aboveground biomass in Central Africa [54,55,52], and used to describe the allometric relationships for a wide range of plants [56]. Power function models also express the allometry between different parts of the plant, such as the proportionality in the relative increments between stem biomass and girth of the trees [6].

## Conclusion

This study provides a new destructive dataset for *Acacia mangium* Willd. plantations on the Batéké Plateau, covering the early growth stages (25–66 months). By integrating 54 destructively sampled trees, power-law models based on D^2^H were identified as the most reliable predictors of tree volume and aboveground biomass. These equations improve the accuracy of biomass and carbon stock estimation compared to default IPCC values and are directly applicable to MRV systems for REDD+ and other forest-based mitigation initiatives. Although the results are constrained by a relatively narrow diameter range (4–24 cm), they provide an essential reference for forest plantations in the Congo Basin and highlight the need for further studies encompassing older stands and larger trees. Overall, this work strengthens the scientific basis for monitoring *A. mangium* Willd. plantations and contributes to more transparent and reliable reporting of forest carbon dynamics.

## Supporting Information

**Figure S1.**
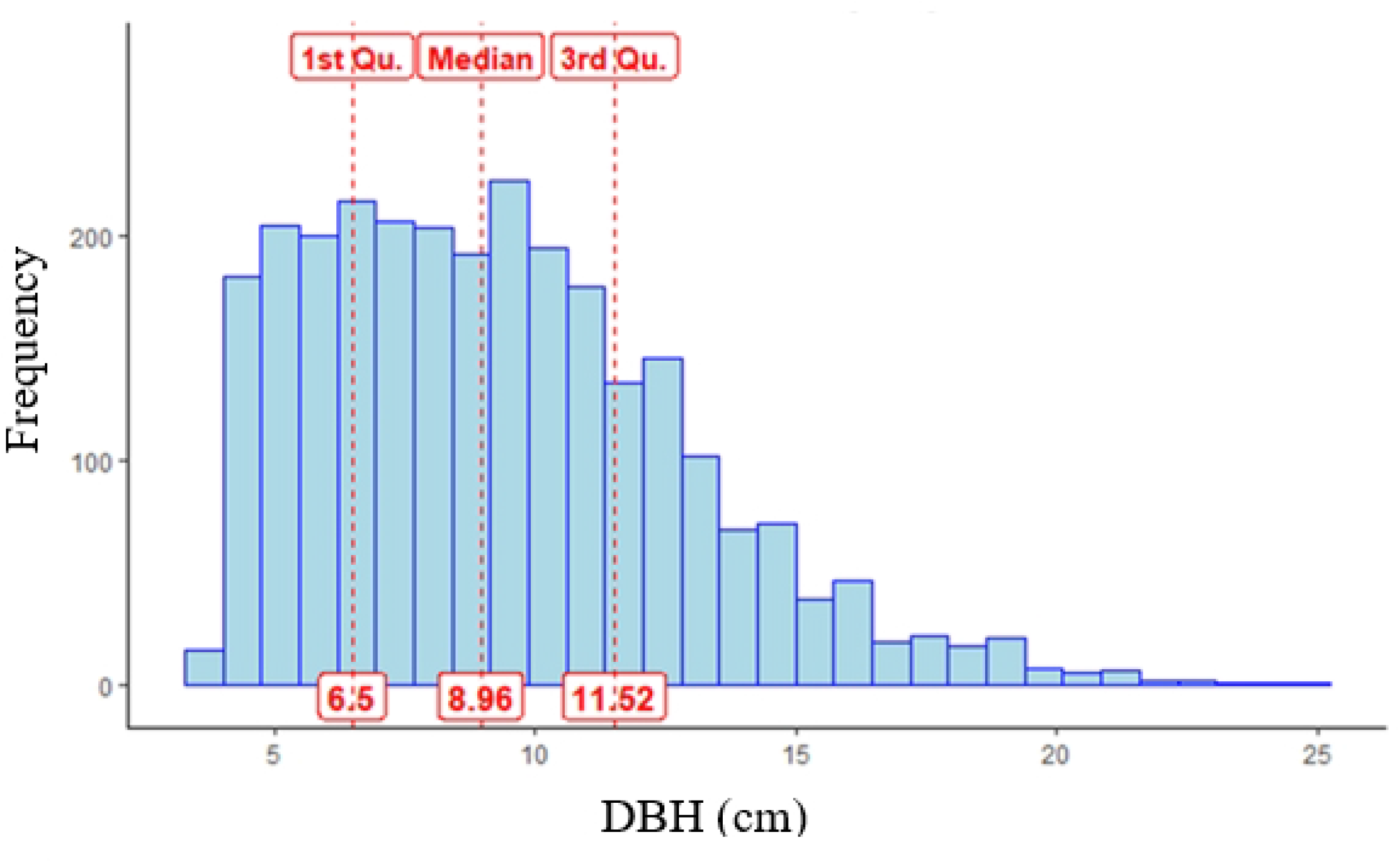
The stand’s DBH distribution of pre-inventory data

**Figure S2.**
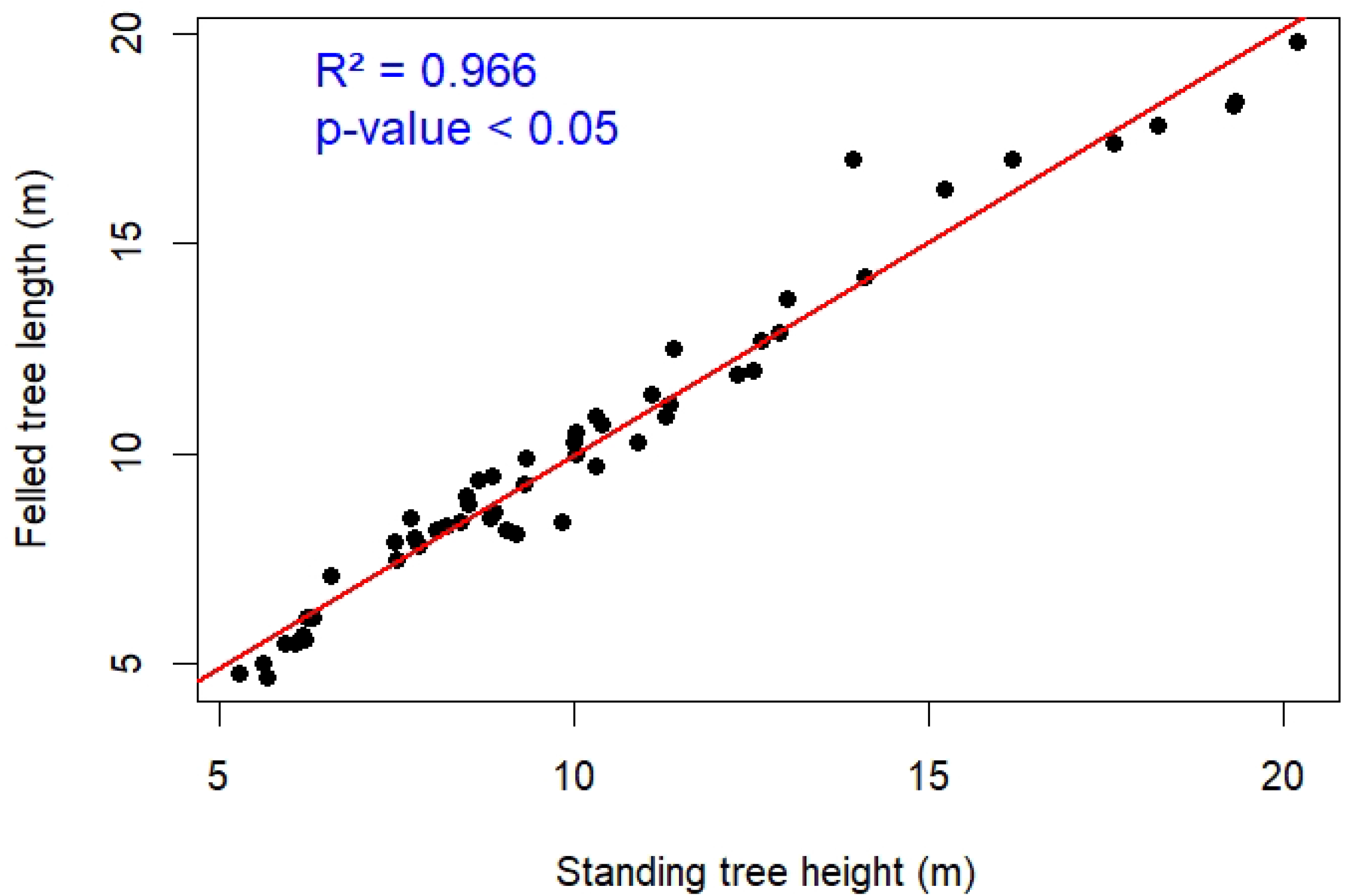
Regression between tree height non-destructive measurement and tree felled length

**Figure S3.**
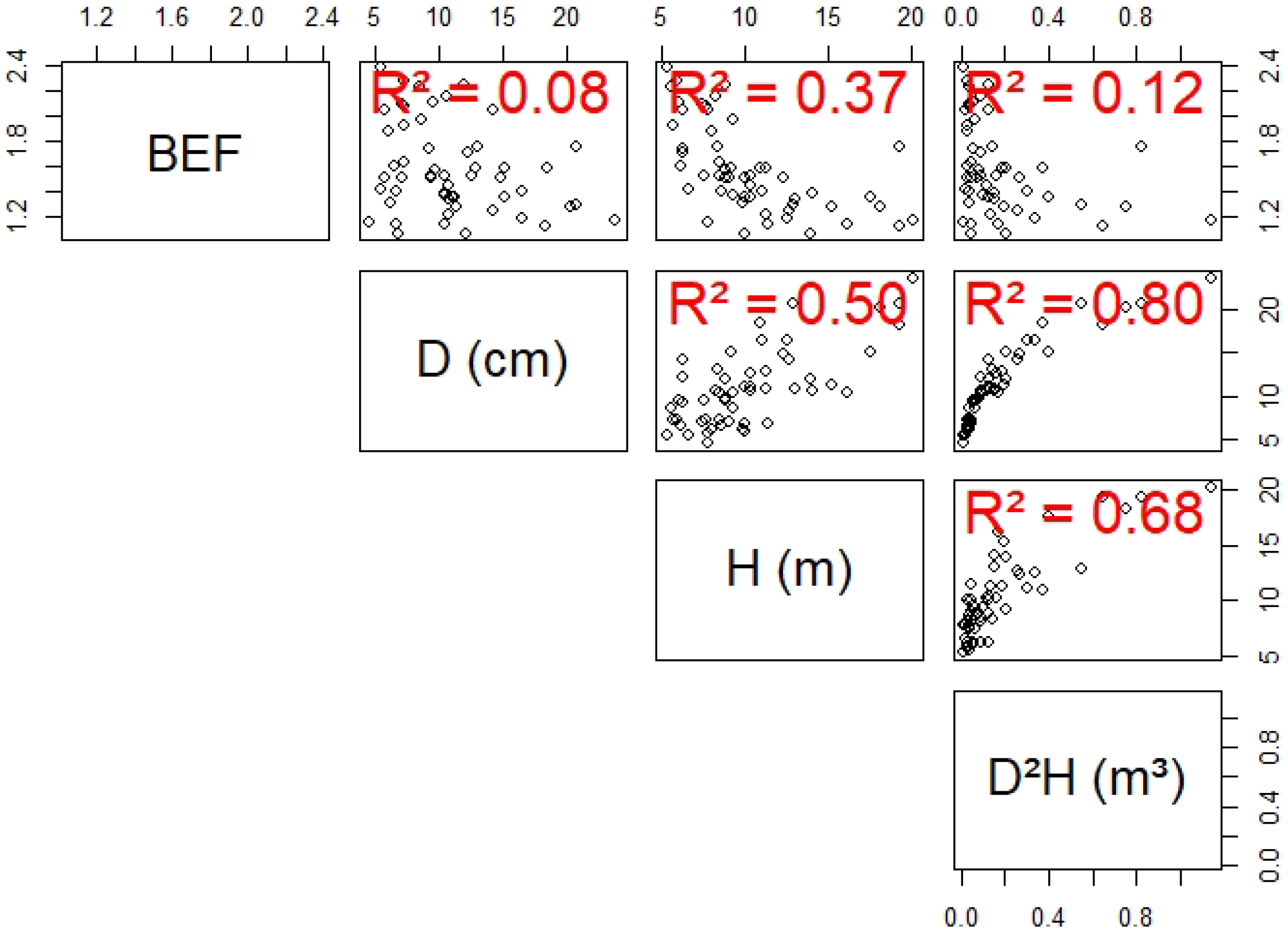
Relationship between BEF and D, H and D^2^H

**Table S1.** Comparison of total tree height measurements methods (repeated measures ANOVA).

| Site | Age<br>(months) | Mean DBH<br>(cm) | Total height (Ht) of stand trees (m) |  |  |  | Total length<br>of felled tree<br>(m) |
| --- | --- | --- | --- | --- | --- | --- | --- |
|  |  |  | *Position 1* | *Position 2* | *Position 3* | *Mean Position* |  |
| Site 1 | 66 | 12.7 ± 5.7 | 14.4 ± 3.5 | 14.4 ± 3.8 | 15.0 ± 4.2 | 14.6 ± 3.8 | 14.7 ± 3.8 |
| Site 2 | 42 | 12.0 ± 4.4 | 10.4 ± 1.7 | 10.3 ± 1.7 | 10.3 ± 1.5 | 10.3 ± 1.6 | 10.4 ± 1.5 |
|  | 25 | 9.9 ± 2.8 | 8.6 ± 1.5 | 8.5 ± 1.3 | 8.1 ± 1.4 | 8.4 ± 1.4 | 8.5 ± 1.1 |
| Site 3 | 25 | 9.7 ± 4.4 | 9.2 ± 1.9 | 9.3 ± 1.8 | 9.2 ± 1.7 | 9.2 ± 1.8 | 9.5 ± 1.5 |
|  | 25 | 9.5 ± 3.3 | 6.2 ± 1.2 | 6.4 ± 1.1 | 6.2 ± 1.0 | 6.3 ± 1.1 | 5.7 ± 1.0 |
| Average |  | 11.0 ± 4.5 | 10.3 ± 3.7 | 10.3 ± 3.7 | 10.4 ± 4.0 | 10.3 ± 3.8 | 10.3 ± 3.9 |
| repeated measures ANOVA |  |  | p-value = 0.911 (> 0.05); F(3,159) = 0.178; ges = 0.000064 |  |  |  |  |

*Raw distributions were non-normal (Shapiro–Wilk, $p \leq 0.001$ for all four methods), but normality was achieved after log transformation ( $p = 0.097–0.219$ ). A repeated-measures ANOVA on log-transformed data showed no method effect ( $F(3,159) = 0.178$ , $p = 0.911$ ; generalized eta squared (ges) = 0.000064. These findings indicate that measures at Position 1, 2, 3, and Felled yield equivalent total height estimates*

## Acknowledgment

The authors wish to express their sincere gratitude to the interns Aziz NABINE, Suveny IKIA, Amazone Pulcherie BOTETI, and Philippe Joseph TIKENG for their contribution to field data collection. Thanks to Jeanne Clémence for her support in supervising the internship team during part of the study. We further extend our appreciation to the entire FRM and SPF2B teams for their logistical and operational support, which greatly contributed to the successful completion of this work.

## Author contributions

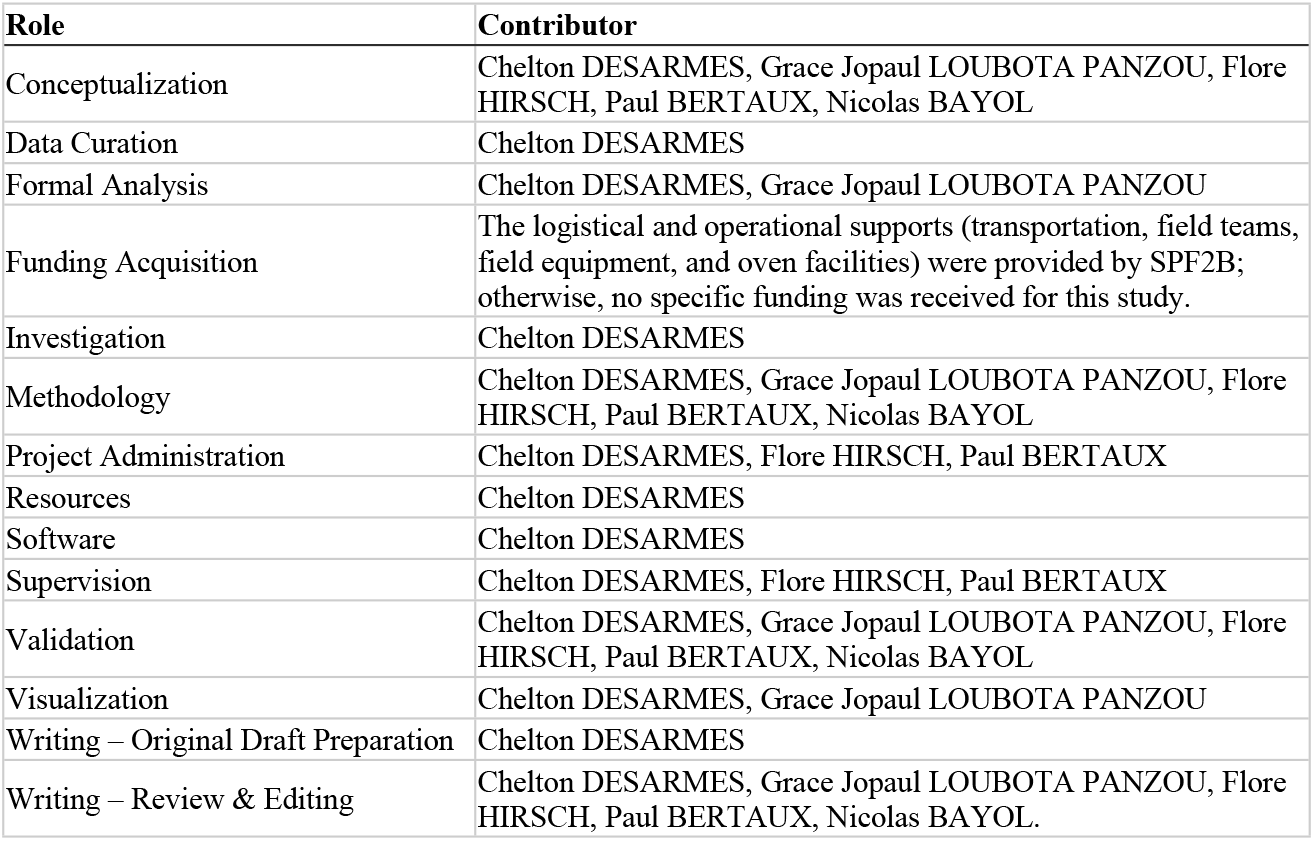

## References

1. Gibbs HK, Brown S, Niles JO, Foley JA. Monitoring and estimating tropical forest carbon stocks: making REDD a reality. Environ Res Lett. 2007;2(4):045023.

2. Pechanec V, Štěrbová L, Purkyt J, Prokopová M, Včeláková R, Cudlín O, et al. Selected Aspects of Carbon Stock Assessment in Aboveground Biomass. Land. 2022;11(1):66.

3. Maniatis D, Scriven J, Jonckheere I, Laughlin J, Todd K. Toward REDD+ Implementation. Annu Rev Environ Resour. 2019;44(1):373–98.

4. Henry M, Picard N, Trotta C, Manlay R, Valentini R, Bernoux M, et al. Estimating Tree Biomass of Sub-Saharan African Forests: a Review of Available Allometric Equations. Silva Fenn. 2011;45(3):477–569.

5. Vashum KT, Jayakumar S. Methods to Estimate Above-Ground Biomass and Carbon Stock in Natural Forests - A Review. J Ecosyst Ecography. 2012;2(4):1–7.

6. Picard N, Saint-Andre L, Henry M. Manuel de construction d’équations allométriques pour l’estimation du volume et la biomasse des arbres. De la mesure de terrain à la prédiction. 2012.

7. Chave J, Réjou-Méchain M, Burquez A, Chidumayo E, Colgan M, Delitti W, et al. Improved allometric models to estimate the aboveground biomass of tropical trees. Glob Change Biol. 2014;20:3177–90.

8. Chave J, Andalo C, Brown S, Cairns MA, Chambers JQ, Eamus D, et al. Tree allometry and improved estimation of carbon stocks and balance in tropical forests. Oecologia. août 2005;145(1):87–99.

9. Alvarez E, Duque A, Saldarriaga J, Cabrera K, De Las Salas G, Del Valle I, et al. Tree above-ground biomass allometries for carbon stocks estimation in the natural forests of Colombia. For Ecol Manag. 2012;267:297–308.

10. Zianis D, Mencuccini M. On simplifying allometric analyses of forest biomass. For Ecol Manag. 2004;187(2-3):311–32.

11. Bastin JF, Finegold Y, Garcia C, Mollicone D, Rezende M, Routh D, et al. The global tree restoration potential. Science. 2019;365(6448):76–9.

12. Norisada M, Hitsuma G, Kuroda K, Yamanoshita T, Masumori M, Tange T, et al. Acacia mangium, a Nurse Tree Candidate for Reforestation on Degraded Sandy Soils in the Malay Peninsula. For Sci. 1 oct 2005;51(5):498–510.

13. Bernhard-Reversat F, Diangana D, Tsatsa M. Biomasse, minéralomasse et productivité en plantation d’Acacia mangium et A. auriculiformis au Congo. BOIS FORETS Trop. 1993;238:35–44.

14. Hoare AL. THE USE OF NON-TIMBER FOREST PRODUCTS IN THE CONGO BASIN: London, UK.: The Rainforest Foundation; 2007 p. 53.

15. FAO. Global Forest Resources Assessment 2020. Rome: FAO; 2020.

16. République du Congo. Contribution Déterminée au Niveau National (CDN). Brazzaville, Congo: Ministère du Tourisme et de l’Environnement,.; 2021 p. p32.

17. MEFDD. MINISTÈRE DE L’ÉCONOMIE FORESTIERE ET DU DEVELOPPEMENT DURABLE. 2014. Programme National d’Afforestation et de Reboisement. Décret n° 2013 - 221 du 30 mai 2013. Disponible sur: https://economie-forestiere.gouv.cg/le-ministere/organismes-sous-tutelle/pronar/

18. National Research Council. Mangium and Other Fast-Growing Acacias for the Humid Tropics. Washington, D.C.: National Academies Press; 1983.

19. Koutika LS. Afforesting savannas with Acacia mangium and eucalyptus improves P availability in Arenosols of the Congolese coastal plains. Geoderma Reg. mars 2019;16:e00207.

20. Benbrahim KF, Berrada H, Ghachtouli NE, Ismaili M. Les acacias: des plantes fixatrices d’azote prometteuses pour le développement durable des zones arides et semi-arides. Int J Innov Appl Stud. 2014;8(1):46–58.

21. Fikri Benbrahim K, Berrada H, el ghachtouli N, Ismaili M. Les acacias: des plantes fixatrices d’azote prometteuses pour le développement durable des zones arides et semiarides [Acacia: Promising Nitrogen fixing trees for sustainable development in arid and semi-arid areas. Int J Innov Appl Stud. x1 sept 2014;8:46–58.

22. Krisnawati H., Kallio M.H., Kanninen M. Acacia mangium Willd.: Ecology, silviculture and productivity. Center for International Forestry Research (CIFOR); 2011.

23. Mayola M, Kabamba C, Komanda J. Croissance et production des peuplements d’Acacia mangiun d’âges différents sur le sol sableux du plateau de Batéké/RD Congo. J OASIS Agric Sustain Dev. 3 mai 2023;5:36–46.

24. Semeki Ngabinzeke J. Rendement des fours traditionnels de production de charbon à Acacia auriculiformis et A. mangium sur le plateau des Batéké, RD Congo. 2024;

25. Boissezon P de. Les sols de savane des plateaux Batéké.

26. Alain LM. Contribution à l’étude des plateaux Batékés (géologie, géomorphologie, hydrogéologie)-fdi:00743-Horizon.

27. Siarudin M, Indrajaya Y. Biomass estimation model for small diameter Auri tree (Acacia auriculiformis A. Cunn. ex Benth.). IOP Conf Ser Earth Environ Sci. 2019;308(1):012028.

28. CTFT. Mémento du forestier tropical. 3e éd. 1989. Paris: Ministère de la coopération; 1989. 1266 p. (Techniques rurales en Afrique).

29. Pardé Jean Bouchon Jean. Dendrométrie. 2e édition refondue. Nancy: École nationale du génie rural, des eaux et des forêts; 1988.

30. Lejeune P, Rondeux J. Modèles de cubage pour essences multi-tiges : application à des plantations d’acacia. Cah Agric [Internet]. 1994;3(3). Disponible sur: https://orbi.uliege.be/handle/2268/24945

31. Peroches A, Deleporte P. Etude sur la productivité en bois de différentes provenances d’Acacia auriculiformis et Acacia mangium sur le Plateau Batéké en République Démocratique du Congo; 2014;

32. Louppe D, M’Bla Koua, Coulibaly A. Tarifs de cubage pour Afzelia africana Smith en forêt de Badénou (Nord Côte d’Ivoire). 1994; Disponible sur: https://agritrop.cirad.fr/311985/

33. Rondeux J. La mesure des arbres et des peuplements forestiers. 2021.

34. Zanne A, Lopez-Gonzalez G, Coomes D, Ilic J, Jansen S, Lewis S, et al. Global wood density database. In 2009.

35. Adam NS, Jusoh I. Allometric Model for Predicting Aboveground Biomass and Carbon Stock of Acacia Plantations in Sarawak, Malaysia. BioResources. 14 août 2018;13(4):7381–94.

36. Brown S, Gillespie AJR, Lugo AE. Biomass Estimation Methods for Tropical Forests with Applications to Forest Inventory Data. For Sci. 1989;35(4):881–902.

37. Malhi Y, Baker TR, Phillips OL, Almeida S, Alvarez E, Arroyo L, et al. The aboveground coarse wood productivity of 104 Neotropical forest plots. Glob Change Biol. 2004;10(5):563–91.

38. Dawkins HC. Estimating total volume of some Caribbean trees. Carib For. 1 janv 1961;22(3/4):62-3.

39. Larjavaara M, Muller-Landau HC. Measuring tree height: a quantitative comparison of two common field methods in a moist tropical forest. Methods Ecol Evol. 2013;4(9):793–801.

40. Hunter MO, Keller M, Victoria D, Morton DC. Tree height and tropical forest biomass estimation. Biogeosciences. 2013;10(12):8385–99.

41. Baskerville GL. Use of Logarithmic Regression in the Estimation of Plant Biomass. Can J For Res. 1972;

42. Sprugel DG. Correcting for Bias in Log-Transformed Allometric Equations. Ecology. 1983;64(1):209–10.

43. Miyakuni K, Heriansyah I, Heriyanto NM, Kiyono Y. Allometric Biomass Equations, Biomass Expansion Factors and Root-to-shoot Ratios of Planted Acacia mangium Willd. Forests in West Java, Indonesia. J For Plan. 2004;10(2):69–76.

44. Akaike H. A new look at the statistical model identification. IEEE Trans Autom Control. 1974;19(6):716–23.

45. Rykiel EJ. Testing ecological models: the meaning of validation. Ecol Model. 1996;90(3):229–44.

46. Hastie T, Tibshirani R, Friedman J. The Elements of Statistical Learning. New York, NY: Springer New York; 2009. (Springer Series in Statistics).

47. Yates LA, Aandahl Z, Richards SA, Brook BW. Cross validation for model selection: A review with examples from ecology. Ecol Monogr. 2023;93(1):e1557.

48. Shumie A, Alemu A, Abebe G, Gemtesa G, Gebremariam Y, Alene Y, et al. Allometric equations for estimation of above- and below-ground biomass of Acacia mearnsii in northwestern Ethiopia. For Sci Technol. 2024;20(3):279–85.

49. IPCC. Good practice guidance for land use, land-use change and forestry /The Intergovernmental Panel on Climate Change. Ed. by Jim Penman. Penman J, éditeur. Hayama, Kanagawa; 2003.

50. Chave J, Condit R, Aguilar S, Hernandez A, Lao S, Perez R. Error propagation and scaling for tropical forest biomass estimates. Malhi Y, Phillips OL, éditeurs. Philos Trans R Soc Lond B Biol Sci. 2004;359(1443):409–20.

51. Molto Q, Rossi V, Blanc L. Error propagation in biomass estimation in tropical forests. Freckleton R, éditeur. Methods Ecol Evol. 2013;4(2):175–83.

52. Fayolle A, Ngomanda A, Mbasi M, Barbier N, Bocko Y, Boyemba F, et al. A regional allometry for the Congo basin forests based on the largest ever destructive sampling. For Ecol Manag. 2018;430:228–40.

53. Ketterings QM, Coe R, Van Noordwijk M, Ambagau’ Y, Palm CA. Reducing uncertainty in the use of allometric biomass equations for predicting above-ground tree biomass in mixed secondary forests. For Ecol Manag. 2001;146(1-3):199–209.

54. Proces P, Dubiez E, Bisiaux F, Péroches A, Fayolle A. Production d’Acacia auriculiformis dans le système agroforestier de Mampu, plateau Batéké, République démocratique du Congo. BOIS FORETS Trop. 2 janv 2018;334:23.

55. Fayolle A, Rondeux J, Doucet JL, Ernst G, Bouissou C, Quevauvillers S, et al. Réviser les tarifs de cubage pour mieux gérer les forêts du Cameroun. BOIS FORETS Trop. 2013;317(317):35.

56. Niklas KJ. Size-dependent Allometry of Tree Height, Diameter and Trunk-taper. Ann Bot. 1 mars 1995;75(3):217–27.

